# Insect Identification with ESEM Imaging and DNA Barcoding for Forensic Use

**DOI:** 10.1101/2025.11.05.686861

**Authors:** Riley B. Hoffman, Nicholas J. Miller, Ashley Hall

## Abstract

Flies are among the first insects to arrive at a corpse to lay eggs. The developing larvae can be used to determine post-mortem interval (PMI) using species specific growth models. The larvae can be difficult to identify due to minute morphological characteristics and rearing larvae into adults is time consuming and may be unsuccessful. We investigated the potential of coupling Environmental Scanning Electron Microscopy (ESEM) and DNA barcoding to enable the acquisition of high resolution images while obtaining DNA in a minimally destructive manner. This study focused on the widespread and forensically relevant species *Chrysomya rufifacies* as a model. Specimens were prepared in three different drying treatments: unaltered, dried, dried and imaged. Following the imaging process, three different DNA extraction methods with different levels of sample destruction were tested. ESEM imaging showed that there were no visual differences between specimens that underwent drying treatments and ones that were not dried. No significant differences in DNA yield were observed for drying treatments and extraction methods post extraction. However, extraction methods demonstrated significant differences in yield of PCR products. Sequenced PCR products aligned with 100% identity to the *C. rufifacies* mitochondrial genome reference sequence. Lower nucleotide identities between the sequenced PCR products and other forensically relevant fly species demonstrated that combining ESEM and DNA barcoding is a promising approach to rapid and reliable identification of insect larvae recovered from forensic samples.

## INTRODUCTION

The development and succession rates of carrion insects are important to forensic science for the creation of post-mortem interval (PMI) (1). Calliphorids, commonly known as blow flies, are some of the first insects to arrive at a corpse to lay eggs. Calliphorids are frequently encountered due to their wide geographic distribution and early colonization, making them forensically significant (2,3).

### PMI and Insects

In the first 24-48 hours after death pathological testing on tissue samples and body fluids can be used to determine the post-mortem interval (PMI). After 72 hours pathological evidence is no longer reliable (4) and entomological evidence becomes important. Forensic entomologists use known succession rates and species specific growth models for larvae under similar environmental conditions to calculate PMI (4). The reliance on species specific growth models to perform PMI calculations requires accurate identification of specimens collected from the crime scene. Forensic entomologists play a critical role in identifying and interpreting the temporal significance of finding a particular species at a crime scene for the creation of an investigatory timeline.

### Morphological Identification and Imaging

Identification of Diptera larvae requires extensive training to recognized minute morphological features that can only be seen through a microscope (Figure 1) (5). Additionally, many of these morphological features only become distinct towards the end of larval development. The larval stage is divided into three instars, with most larval identification keys only being created for the use on the third instar. As an example, third instars of *Chrysomya rufifacies*, a widespread species commonly found colonizing corpses can be identified by small morphological features including the pattern of the protruding tubules along the body (Figure 1C) and the posterior end of the larvae (Figure 1E) that have three rows of spines at the end of each tubule (Figure 1D). Some morphological characteristics may not be species-specific but are used to indicate the third instar, such as three slits on the posterior spiracles (Figure 1F), ten papillae on the anterior spiracle (Figure 1B), and intersegmental spines with 2-4 points (Figure 1A). Due to the specialized nature of the larval identification the Organization of Scientific Area Committees for Forensic Science (OSAC) recommends that both preserved larvae and live larvae be collected from a crime scene, the latter to be reared into adults for easier identification (6). The suggestion to collect live larvae with the intent to rear is based on the fact that species-level identification of larvae is difficult and requires a considerable amount of time and training compared to identification of adult flies (7). However, rearing adults from larvae can be unsuccessful and requires the right environment for the larvae to develop (8). Taking high resolution images of larvae with a Scanning Electron Microscope (SEM) paired with DNA barcoding would save investigators time by reducing the need for rearing larvae into adults.

**Figure 1.**
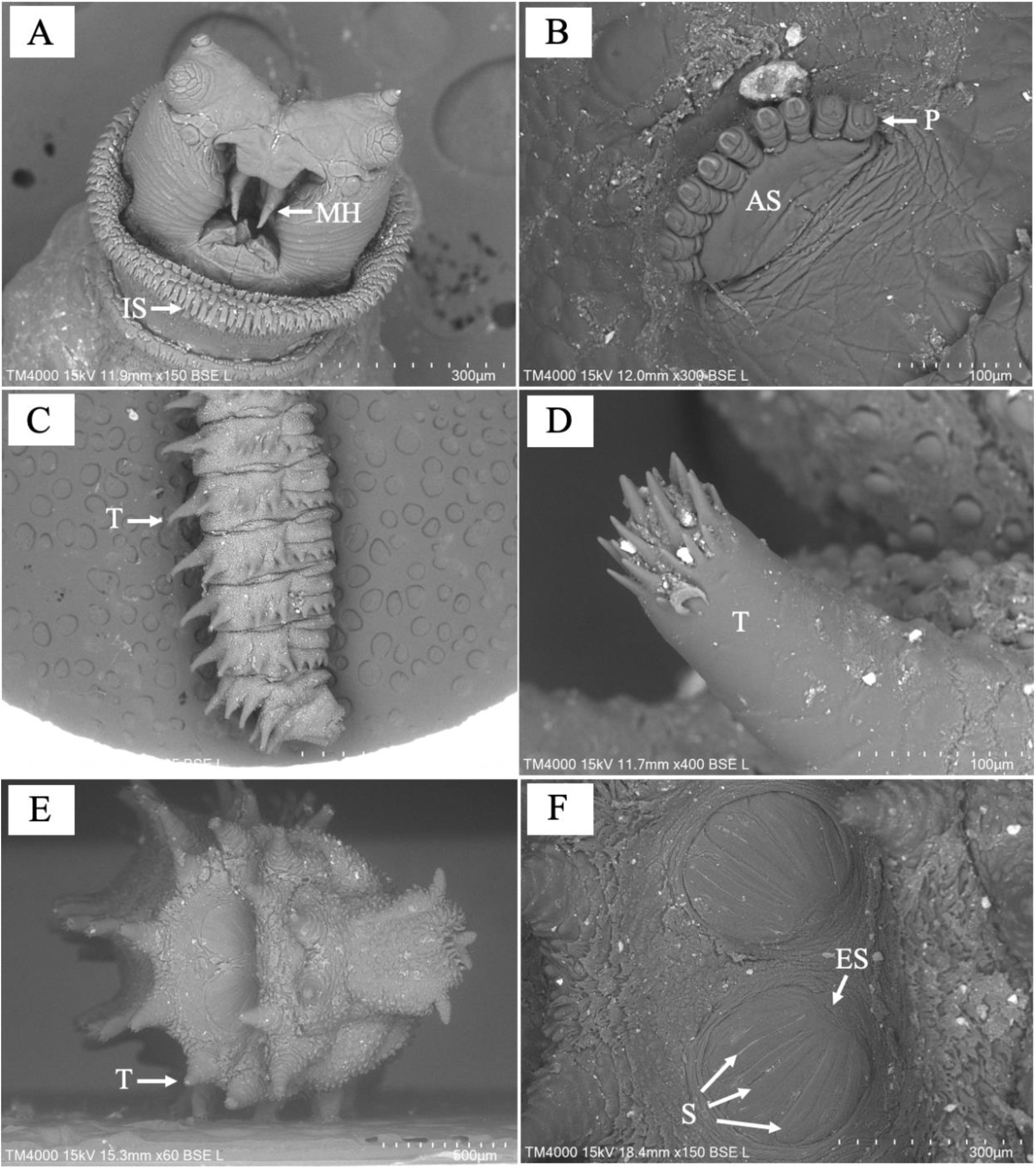
Morphological features used for larval identification as shown on 3^rd^ instar *Chrysomya rufifacies using ESEM*. (A) Anterior end of larva showing prominent mouth hooks (MH) and intersegmental spines (IS) that have 2-4 points on the ends; (B) a fan-like anterior spiracle (AS) with 10 papillae along the edge of the spiracle; (C) elongated tubercles (T) encircling each segment of the larva; (D) three rows of spines present on the end of each elongated tubercle; (E) elongated tubercle (T) surrounding the posterior spiracle; (F) two posterior spiracles with the splits (S) with an ecdysial scar (ES) below the three slits.

Scanning Electron Microscopy (SEM) is considered the gold standard for morphological identification for creating high-resolution images. SEM studies like Sukontason et al. (8) are used by forensic entomologists to compare specimens at all stages of development to known images of specimens to identify the unknown specimens. These studies are important for recording morphological structures across all life stages to develop taxonomic identification keys to characterize stage specific morphological distinctions (8). However, a highly trained individual would be needed to evaluate these minute characteristics for identification past genus level. Additionally, it is more common for Environmental Scanning Electron Microscopes (ESEMs) to be found in an academic entomology lab used for morphological studies since the ESEMs are able to maintain the specimens without complete destruction or major alteration.

### DNA Barcoding

DNA barcoding has become more accessible, making its use to identify insect taxa more common as a complementary technique to SEM imaging. DNA barcoding aims to use short sequences with significant species-level genetic variability for sequence comparisons to reference libraries for species identification (9). The mitochondrial gene cytochrome oxidase I (COI) is often used as the species specific identifier (10).

Pimsler et al. (11) and Kumar et al. (12) showed proof-of-concept for coupling SEM and DNA barcoding, illustrating that it is possible to extract DNA from specimens that have been prepared for SEM imaging. For SEM imaging both Pimsler et al. (11) and Kumar et al. (12) had to coat the specimens in gold to image them with an SEM, requiring the specimen to homogenized for DNA analysis and effectively destroying the specimen. Coupling the tests could make identification of larvae and creating investigatory timelines based on entomological evidence more accessible to forensic investigations. Due to the specialized nature of forensic entomology, this is a skill typically practiced by an expert consultant outside of the crime lab. DNA barcoding could be used to quickly make an identification past genus level and negate the need for rearing. Entomologists can review the images to visually identify the larvae and explain the importance of finding the specimen by estimating PMI.

This procedure potentially reduces timelines and the need for crime scene investigators to collect live specimens to rear to adulthood. By having specialists conducting these tests with an ESEM in a minimally destructive manner, the evidence remains intact and available for cross-examination by other expert witnesses if the case goes to trial. Prior to ESEMs, existing work used SEMs with protocols that required the destruction of the specimen for DNA analysis after imaging. ESEMs are more common within the field of entomology because the specimens require little to no preparation before imaging and keep the insect intact for further analysis and preservation. Allowing further preservation of the specimens also allows for the defense to have their own entomologist to review the morphology of the specimens later as visual identification is still the gold standard.

## MATERIALS AND METHODS

### Sample Collection and Preparation

*Chrysomya rufifacies* were selected as a model organism for this study because they are forensically significant as they are some of the first insects to arrive on a body and have broad geographic distribution, they are easy to rear and were readily available.

Third instar *Chrysomya rufifacies* larvae were collected from a colony kept by Dr. Aaron Tarone’s laboratory at Texas A&M University and preserved in 70% ethanol. Ninety larvae were divided into groups of three drying treatments: unaltered, dried, and dried and imaged.

Unaltered larvae were blotted dry with a Kimwipe® to remove excess ethanol. The dried larvae were placed in a 1:1 solution made of 100% ethanol and 100% acetone, which was then diluted to 70%, 80%, and 90% with DI water. Larvae were immersed in each dilution for 15 minutes moving from the 70% dilution to the 100% dilution.

### ESEM Imaging

The dried and imaged larvae were dried using the same protocol before being imaged using a Hitachi Tabletop Microscope TM4000 and associated software in Dr. Jason Bond’s lab at UC Davis. A piece of double-sided carbon tape was placed on a circular metal stage, the stage was then screwed into a variable tilt mount that could be adjusted to any angle between 0° and 90°. The combined stage and mount was placed in the ESEM where the one to two larvae could be placed on the tape. The chamber was then closed and the TM4000 Software was opened on the connected desktop computer to the user interface page, the ESEM chamber was pressurized. Initial images were received by the software at X25, the controls of the ESEM were used to center the larva on the image receiving screen. Once centered, magnification was increased, the “auto” and “slow” options were selected on the computer software to make the image clearer. The brightness of the image was then adjusted before saving the image and raw data. Images were taken of the posterior and anterior spiracles of each larva.

### DNA Extraction Treatments

The three sample preparation groups were further divided into three extraction treatment groups. Ten larvae from each of the sample preparation groups were left whole, ten had the body perforated by making a small whole with a fine insect pin, and ten where homogenized. All larvae were left overnight to incubate in lysis buffer.

DNA extractions were preformed using the Zymo Research Quick DNA− Miniprep Plus Kit (Zymo Research Operations, Tuslin, CA), following the Solid Tissue Protocols using 50 µL of elution buffer. Extracted DNA was quantified using Qubit− 4 Fluorometer (Thermo Fisher Scientific, Waltham, MA) with the dsDNA High Sensitivity Assay using 5 µL of the eluted DNA.

### PCR Amplification and Gel Visualization

Primers TY-J-1460 (TACAATTTATCGCCTAAACTTCAGCC) and C1-N-2329 (ACTGTAAATATATGATGAGCTCA) were selected from Wells and Sperling (1999) and aligned with a published *Chrysomya rufifacies* mitochondrial genome (accession number NC_019634.1). These primers were aligned in silico to determine the predicted amplicon and its length that we expected to see measured within agarose gels. These primers were selected from a screening of different COI primers because they reliably amplified the predicted amplicon. The PCR amplification mixtures contained the following components: 2 µM primer, 1X Buffer, 1.5 mM of MgCl_2_, 2.5 mM dNTPs, and 1 unit of Taq DNA polymerase per reaction. The volumes of template DNA and nuclease-free water were adjusted so every reaction contained 2.5 ng of template DNA, based on the extraction quantification results. Cycling conditions consisted of initial denaturation of 95°C for 1 min, followed by 35 cycles of denaturation at 95**°**C for 1 min; annealing at 50°C for 1 min; and extension at 72°C for 1 min, and final extension at 72°C for 5 min.

PCR products were visualized in 1.2% agarose gels with 15 µl of each sample and 3µl of Thermo Scientific 6X TriTrack DNA Loading Dye prior to electrophoresis, which were run for 1 hour at 55 volts. The PCR products were compared to the Thermo Scientific GeneRuler DNA size standard (100 bp – 1000 bp). The PCR products were diluted in a 1:10 dilution before being quantified using Qubit− 4 Fluorometer with Qubit− dsDNA Quantification Assay Kit.

### Sequencing and In Silico Analysis

DNA samples with strong amplification yields were selected for re-amplification and sequencing from each of the nine categories. Samples were amplified and quantified again before being purified using the Qiagen MinElute PCR Purification Kit following the MinElute PCR Purification Kit using a Microcentrifuge Protocol using 10 µL of Buffer EB. The eluted products were sent to Psomagen (Rockville, MD) for sequencing.

The sequencing results were cropped using 4Peaks (Nucleobytes, Aalsmeer, Netherlands) to remove the opposite primer and the ends of the sequences that were low-quality. The cleaned forward and reverse sequences were copied into one text file and combined in Galaxy (13) using the cap3 tool (14) to create a single contig. The output sequence was entered into BLAST as a nucleotide sequence to confirm the amplicon was identified as being from *Chrysomya rufifacies* (15). Once all of the sample sequences were assembled, nine other Diptera sequences (Table 1) were obtained from Genbank and were aligned with the amplicon fragment using the “EMBOSS needle” tool in Galaxy (16–18). The other Diptera sequences were trimmed to the same amplicon length so fragments could be evaluated for genus species level identification. The sample sequences and the sequences from the other Diptera were aligned using “ClustalW” (19), the output was then which was entered into Ident and Sim to create a matrix of percent identity (20).

**Table 1.** List of Diptera species used for comparison of percent identity to *Chrysomya rufifacies*.

| Family | Genus | Species | Accession Numbers |
| --- | --- | --- | --- |
| Calliphoridae | *Chrysomya* | *C. rufifacies* | NC_019634.1 |
| Calliphoridae | *Chrysomya* | *C. albiceps* | NC_019631.1:1-15237 |
| Calliphoridae | *Chrysomya* | *C. putoria* | NC_002697.1:1-15004 |
| Calliphoridae | *Chrysomya* | *C. megacephala* | KT272775.1:1-15050 |
| Sacrophagidae | *Sarcophaga* | *S. albiceps* | NC_028413.1:1-14866 |
| Sacrophagidae | *Ravinia* | *R. pernix* | NC_026196.1:1-14882 |
| Sacrophagidae | *Oxysarcodexia* | *O. thornax* | NC_041072.1:1-14885 |
| Muscidae | *Musca* | *M. domestica* | KT272857.1:8-14887 |
| Muscidae | *Haematobia* | *H. irritans* | NC_007102.1:1-14888 |

## RESULTS

### ESEM Imaging

Thirty larvae were dried and imaged before proceeding to DNA extraction. One larva was taken directly out of ethanol and patted dry with a Kimwipe® to be used as a comparison to the larvae that had undergone the full drying treatment. The larvae that had been dried showed no visible changes to the external morphology used for identification such as the anterior and posterior spiracles, and the overall topographical surface of the specimens (SF2). No change in appearance of the samples allows for correct identification from the images and the storage of the specimens with the option of further visual observations at later time.

### DNA Extraction

DNA was successfully recovered and quantified from all samples under all the drying and extraction methods. Once all samples were quantified by fluorometry the results were analyzed with an analysis of variance (ANOVA) using RStudio and visualized with a box plot (Figure 2). Neither the drying treatment nor extraction methods had a significant effect on DNA yield (*p* ≥ 0.05). Since there was no significant difference in DNA yield all samples were used as templates for PCR amplification.

**Figure 2.**
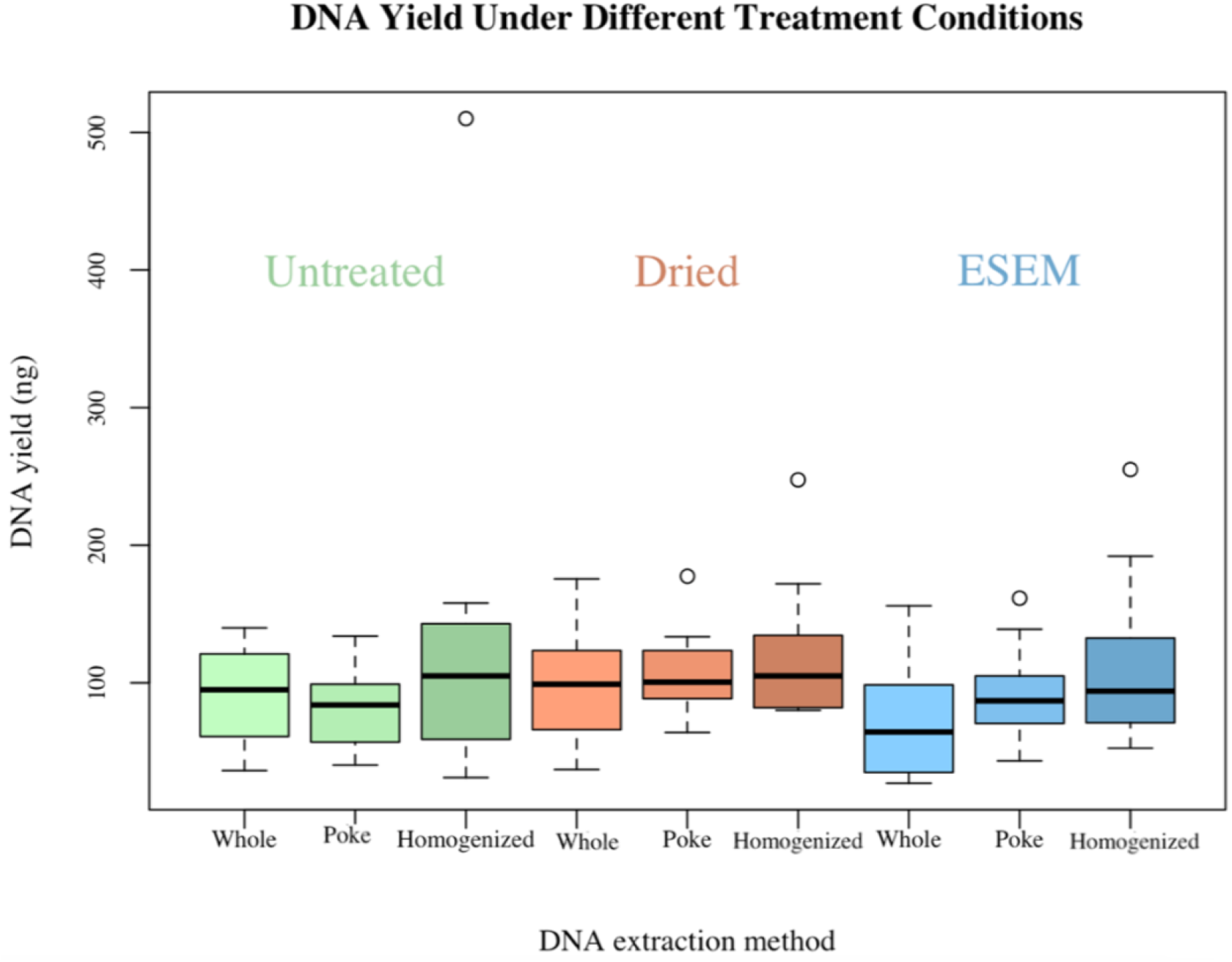
Box plot of DNA yield (ng) per specimen post extraction under different DNA extraction methods and specimen treatments.

### Amplification

Once samples were amplified, they were visualized in an agarose gel to confirm the amplification of the selected fragment based on expected base pair size. All 90 samples were amplified the predicted fragment size. The remaining PCR products were diluted 1:10 with DI water and quantified by fluorometry. An ANOVA was conducted in RStudio to evaluate the effects of extraction method, specimen treatment, and their interaction on DNA concentration after PCR (Figure 3). Extraction method was the only factor to have a significant effect on amplicon yield with a p-value of 2 × 10^−16^. There is a repeating pattern of increasing amplicon yield based on extraction treatment illustrated in Figure 3.

**Figure 3.**
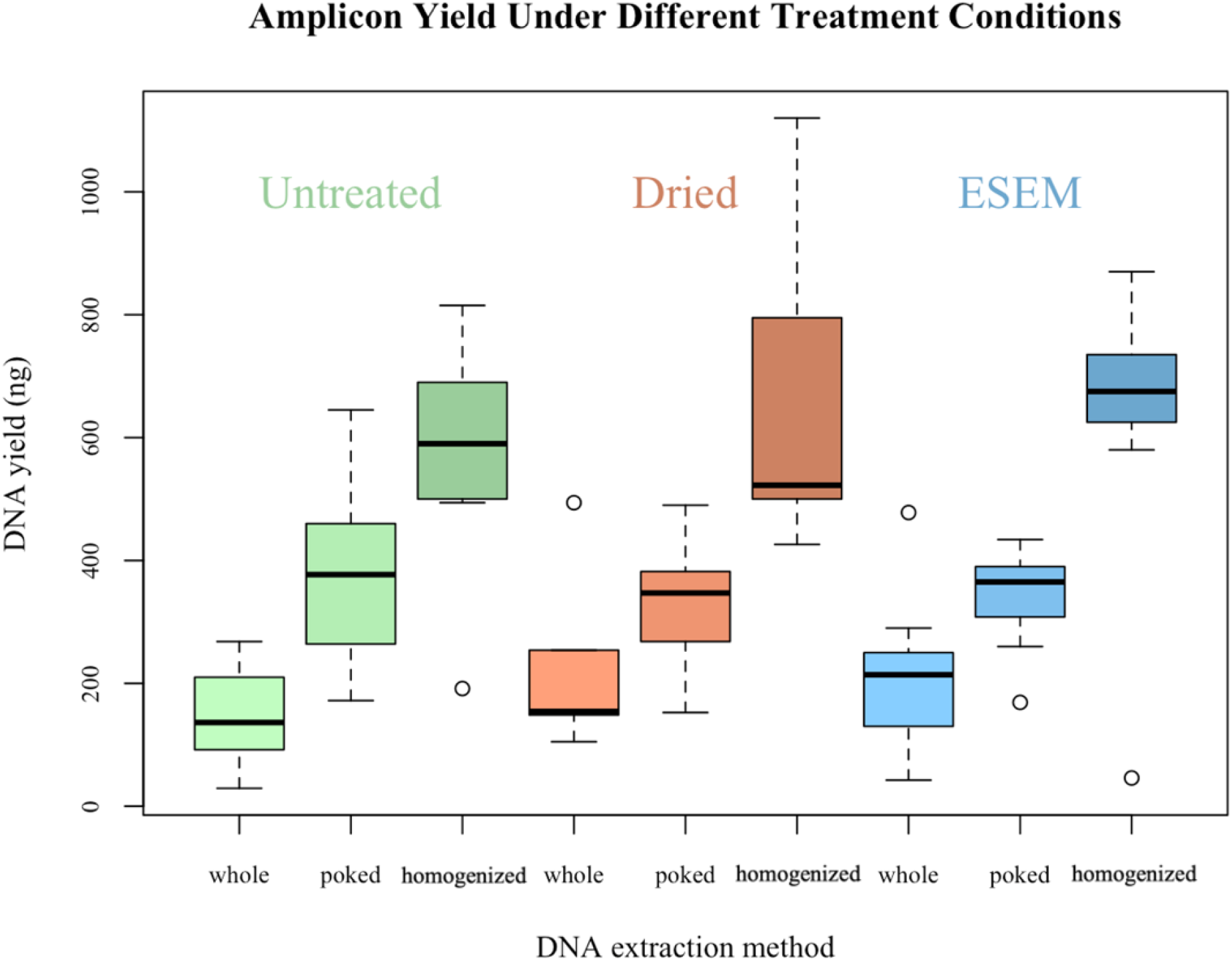
Box plot of amplicon yield (ng) per specimen post amplification under different DNA extraction methods and specimen treatments.

### Sequencing

Sequencing is used to confirm the identity of the amplified fragments which were initially observed for the expected length in agarose gels. Sequencing data was generated for six samples, two of each extraction method – whole, poked, and homogenized – under all drying treatments – unaltered, dried, and dried and imaged. The assembled fragments from the six samples were aligned with sequences of various Diptera from the families Sarcophagidae, Calliphoridae, and Muscidae (Table 1). The aligned sequences were used to compute the similarities of the sequences, visualized in a pairwise matrix of percent identity (Table 2). All six sample sequences had 100% identity to the *Chrysomya rufifacies* reference sequence, regardless of drying treatment or extraction method the samples. Other Diptera sequences showed lower similarity percentages, demonstrating that relationships within the same family and outside of the family can be identified and separated when performing these methods for species identification.

**Table 2.**
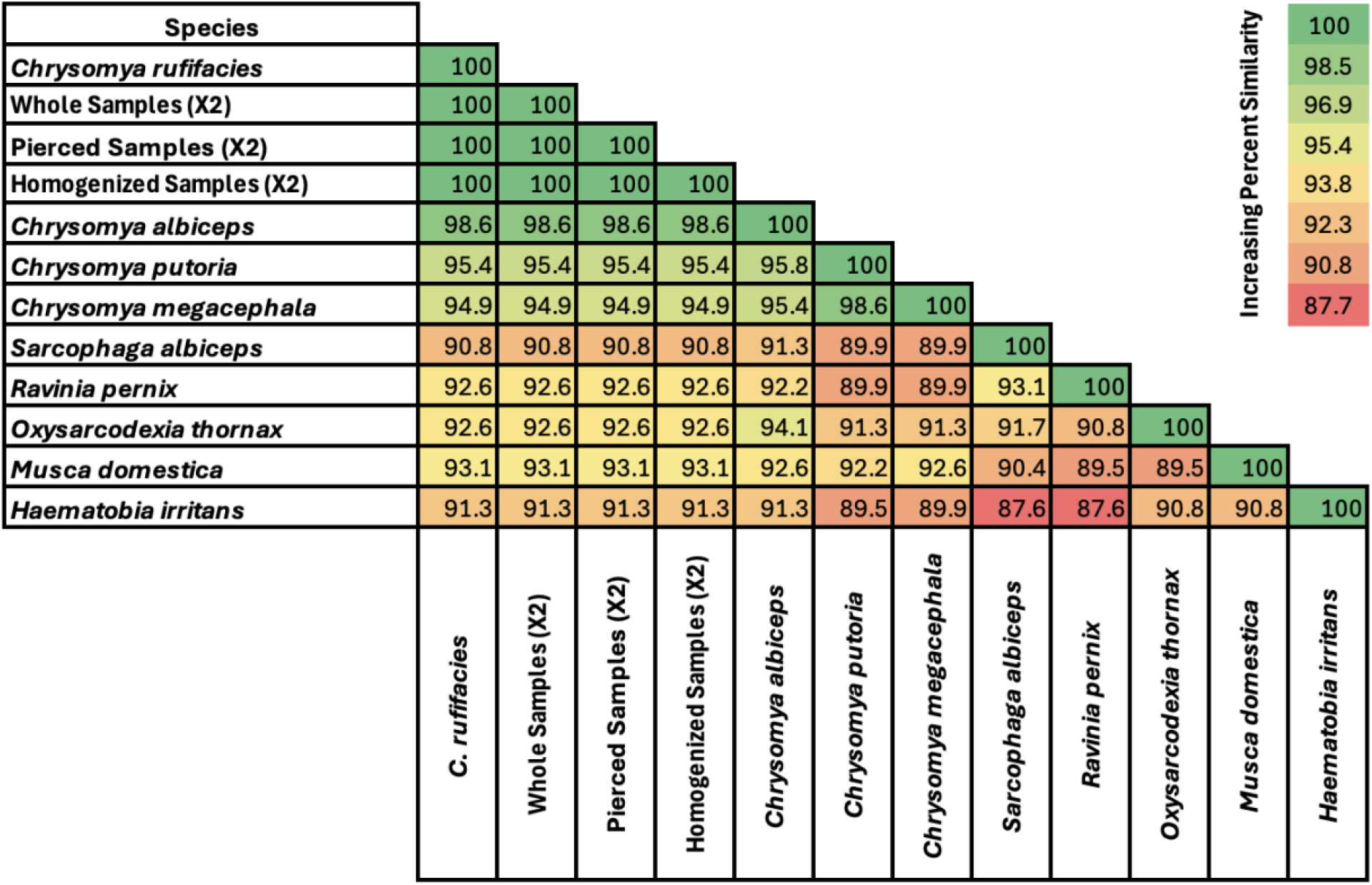
Pairwise matrix of percent identity across six samples and nine reference sequences from other Diptera.

## DISCUSSION

### Imaging

ESEM imaging is the gold standard for high resolution microscopy of morphological structures for identification. Observations across all larval instars of *Chrysomya rufifacies* have been recorded to identify distinguishing features and morphological changes in external structures larvae (8). For our experiments, one larva was taken out of ethanol, blotted dry and imaged for comparison with the larvae that had gone through the drying process. No major differences were noted between the larva directly removed from ethanol and imaged and the specimens that has been dried before imaging (S2). Only third instar specimens were examined because they have the most differentiating external morphology and are easily evaluated as a test species. All dried larvae were successfully imaged with morphological structures unchanged and usable for identification.

ESEMs use a lower vacuum, which allows for higher vapor pressure in the chamber, meaning that it is not necessary for specimens to be dried prior to imaging (21). Drying protocols used in this study closely followed published protocols for SEM imaging, which require specimens to be completely dried before imaging While drying specimens for ESEM imaging is not required, undried specimens lead to increased vapors present in the chamber of the ESEM may contribute to more noise – blurriness (21). While drying is not required, it allows for sharper images even though it could alter morphology. This research was a preliminary study which focused on demonstrating successful protocols for third instar larvae, it remains to be seen what protocols would be necessary for earlier instars.

### DNA Extraction

The *Chrysomya rufifacies* in this study were stored in 70% ethanol for four to five months before being dried for imaging, followed by minimally destructive DNA extraction methods typically used for museum specimens. It has been shown that DNA can be obtained in a minimally destructive manner from larval specimens stored in ethanol for an extended period of time (5,22).

DNA yield for the extraction method did not show any significant difference between the treatments used on the samples (Figure 2). Whereas the quantification of amplicons showed a significant difference in extraction methods (Figure 3). We had expected that both ANOVAs would show the same protocols as significant between the extraction and amplification quantification. One possibility for only the amplification quantification showing that the extraction method was significant was the use of the Qubit™ 4 Fluorometer for the quantification of all extracted samples and post-PCR samples. The Qubit assay is highly specific for detecting double-stranded DNA and measures total double-stranded DNA present in a sample, not exclusively Diptera DNA. Since Qubit™ quantification is not specific to the COI gene, it is possible DNA from the larvae’s blood meals and contaminating microorganisms was also being quantified (23,24). Given the quantification method, it is unknown how much Diptera DNA was added to the PCR mixtures when 2.5 ng of extracted DNA was added. If less Diptera DNA was added to the PCR mixtures this could have contributed to the amplicon quantification trends because there would have been less Diptera DNA to amplify if it was diluted down by other DNA. To improve the success of PCR, nested PCR could be used in future works to refine larger amplicon regions, increase amplicon yield, and improving identification accuracy for DNA barcoding (25).

It is also possible that the DNA quantified after extraction was damaged, causing significant differences in the amplicon quantification. Single-strand breaks and double-strand breaks may occur from internal or external factors to the cell, with single stranded breaks being more common (26,27). Depending on the type of DNA lesions, it can negatively affect amplification efficiency of the selected sequence (28). In the future the use of assay to detect the extent of DNA damage paired with qPCR could be done to more accurately calculate the undamaged DNA to ultimately increase amplification efficiency (28). Similarly, qPCR and specific primers could be quantifying DNA from the blood meals and contaminating microorganisms extracted. However, qPCR only identifies DNA targeted by the assay primers. To fully identify the contents of the samples high throughput sequencing would be necessary.

Another variable that may have contributed to the amount of DNA seen after extraction but not in the amplification could be the presence of extracellular DNA. Insects have an open circulatory system with hemolymph freely moving within an open body cavity. Double-stranded, extracellular DNA found on and within the larva may have increased the quantitative values of the extracted DNA but were not part of the template DNA region used during amplification (29,30).

### DNA barcoding

Six sequences were trimmed and assembled, showing 100% identity to a databased *Chrysomya rufifacies* sequence (Table 2). Utilizing public databases for reference genomes is important for identification via sequencing to compare the variability within and between species. The difference of percent identity for distinguishing *Chrysomya rufifacies* from congenerics ranged from 1.4-5.1%. While other studies have suggested the need for a minimum 3% threshold between species to show the delineation between species and being a high enough threshold to account for sequencing error (31–33). However, it has also been found that interspecific variation between *Chrysomya* species can be less than 0.5%, which suggests the need for the use of the additional gene, ITS2, to reliably identify *Chrysomya* to species (34). Additionally, Yusseff and Agnarsson (35) assert that the COI gene does not reliably distinguish between closely related species of Calliphorids. While *Chrysomya rufifacies* has yet to be reported as one of the species that cannot be distinguished using COI alone, further investigation of the variability of the COI gene in wild-caught *Chrysomya rufifacies* would clarify the power of COI sequences to identify to species level.

Given the genetic similarities between the *Chrysomya* species, an entomologist may have to rely on their knowledge of the local species based on geographical locations and temporal resource partitioning. While two *Chrysomya* species may be located in the same region, the time of year they are most active may differ (36). The insects’ activity during certain times may also indicate a preference for certain temperatures for optimal development (36). Given that PMI calculations are dependent on temperature, optimal temperatures for development are species specific, making growth models for sister species nontransferable (36). Knowing that the development of *Chrysomya* sister species can differ enough to effect PMI calculations, correct identification of larvae is paramount.

## CONCLUSION

Coupling ESEM and DNA barcoding can be used to quickly identify specimens to species while maintaining the specimen for future examination. The methodologies used are best preformed in an academic entomology lab because of the access to ESEM instruments.

We have demonstrated that DNA barcoding can successfully identify larvae regardless of treatment and extraction method. Coupling ESEM and DNA barcoding makes forensic entomology more accessible in a shorter timeframe by reducing the need for rearing out adult flies. This study has shown the potential of this methodology which should be expanded to include first and second instars to find the best practices for each instar of *Chrysomya rufifacies*. Once reliable methods for each instar have been optimized, a nationwide sample set of wild *Chrysomya rufifacies* should be created to study the variability of this segment of COI.

## Supporting information

Supplemental images

## SUPPLEMENTAL MATERIAL

**Supplemental Figure 1.**
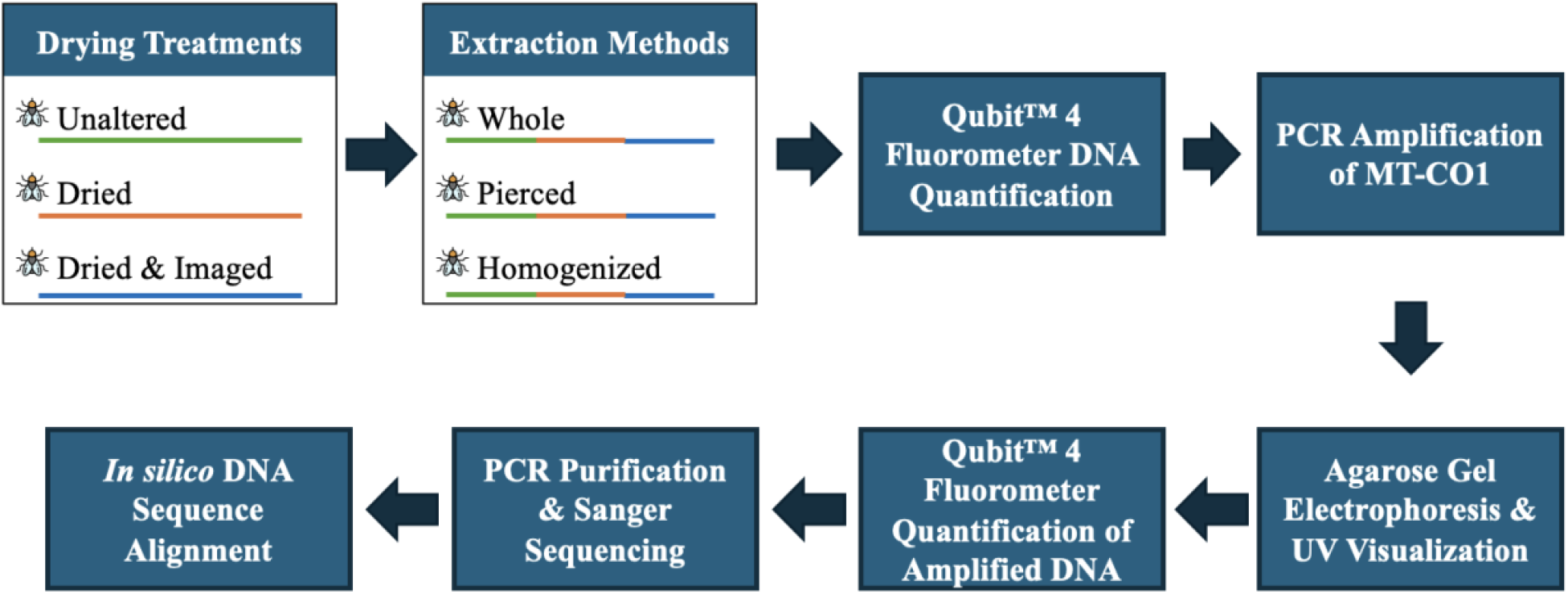
Overview of the experimental workflow, from specimen treatment DNA extraction to PCR amplification, sequencing, ending at data analysis.

**Supplemental Figure 2.**
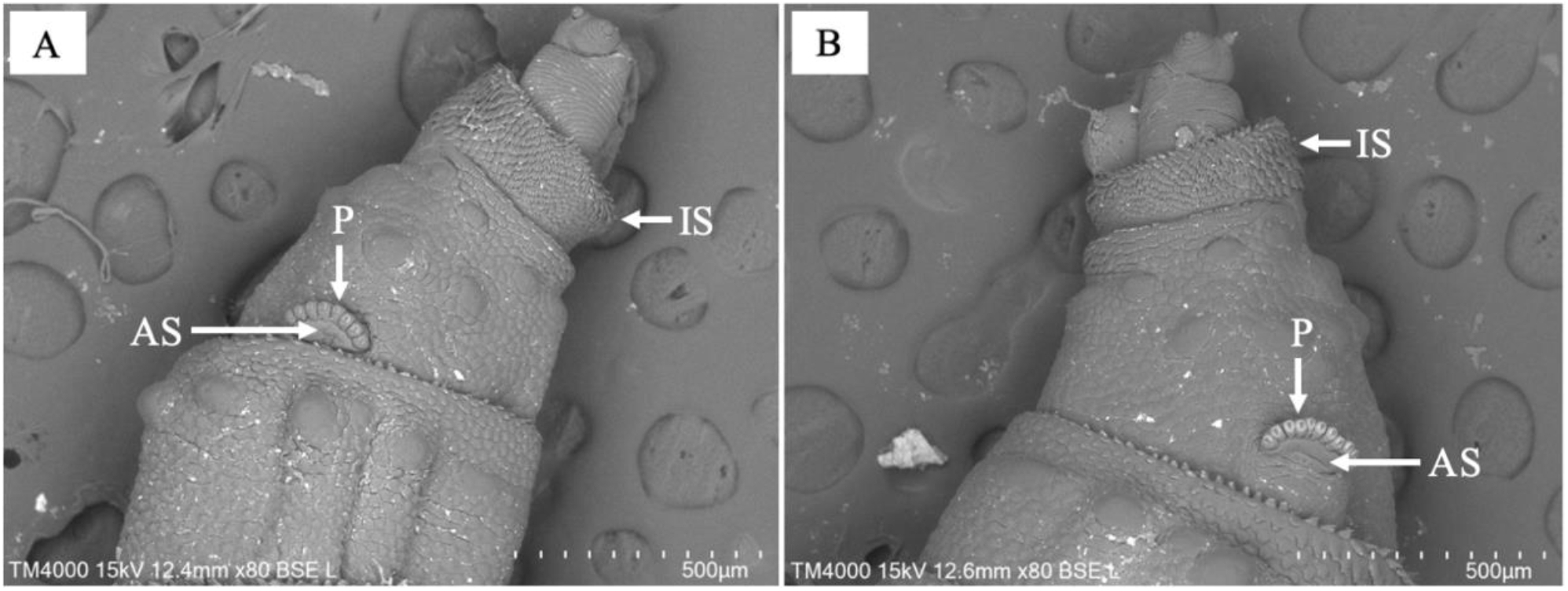
Comparison of the anterior end of Chrysomya rufifacies larvae using ESEM at 80x magnification with BSE detection signal in left orientation: (A) dried larva sample and (B) undried larva sample.

**Supplemental Table 1.**
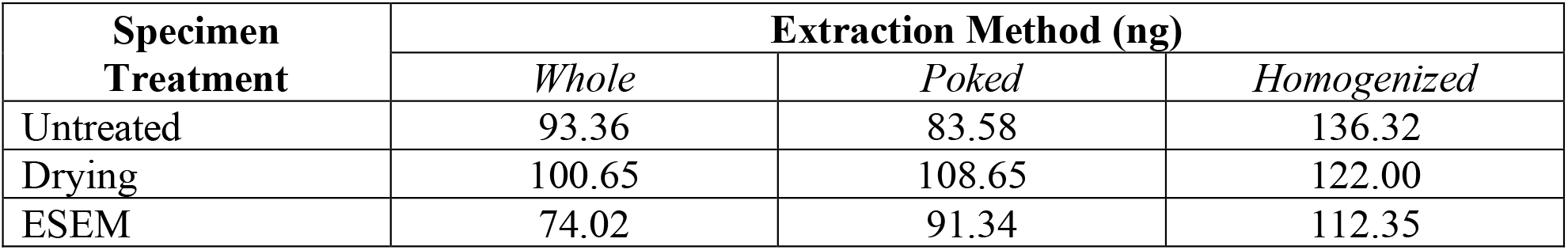
Total DNA Yield (ng) post DNA extraction from different specimen treatments per extraction method.

**Supplemental Table 2.**
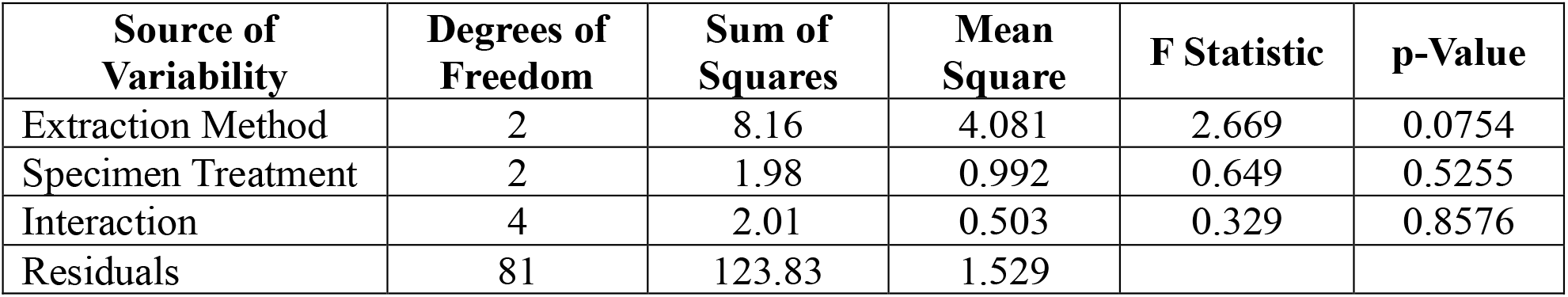
Analysis of Variance (ANOVA) of DNA yield (ng/µL) from larval specimens post extraction using different extraction methods and specimen treatments.

**Supplemental Table 2.**
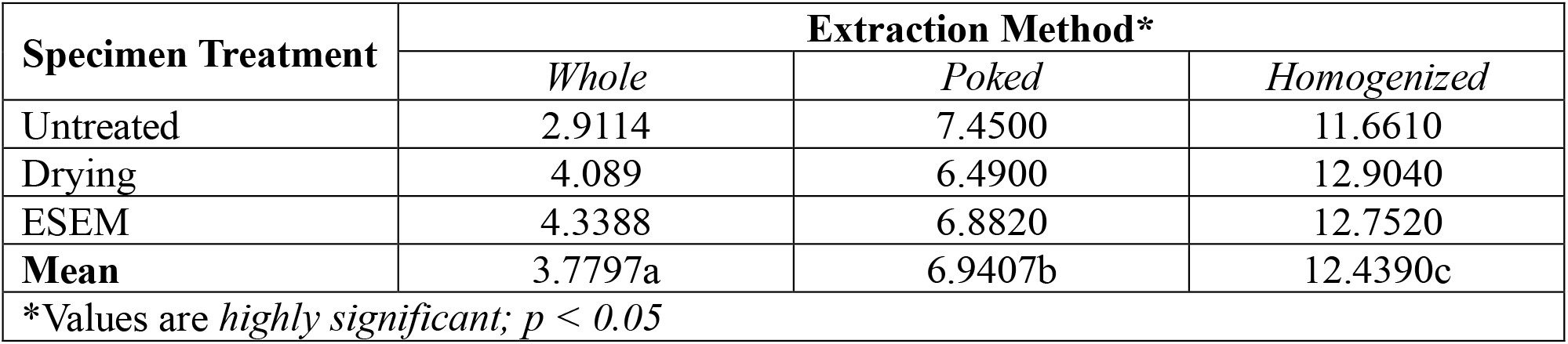
Mean of amplicon concentration (ng/µL) post PCR from different specimen treatments per extraction method.

**Supplemental Table 3.**
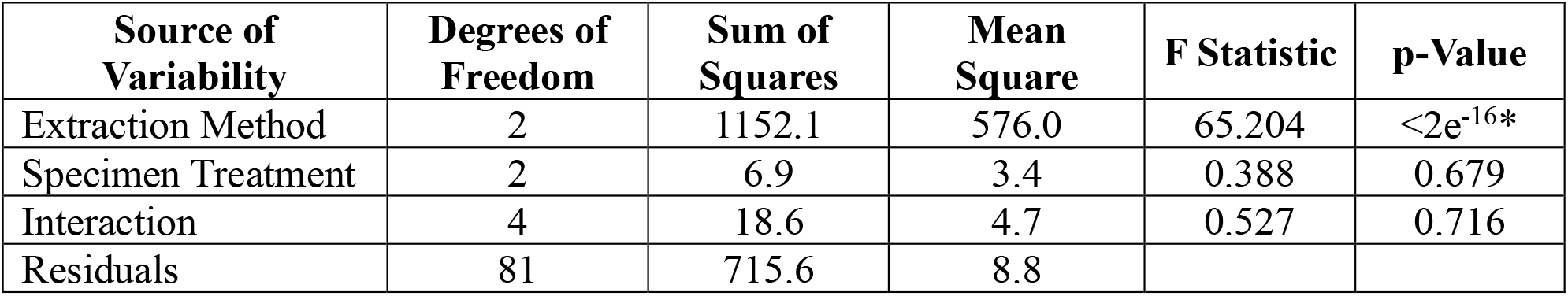
Analysis of Variance (ANOVA) of DNA concentration (ng/µL) from larval specimens post PCR using different extraction methods and specimen treatments.

## REFERENCES

1. Matuszewski S. Post-Mortem Interval Estimation Based on Insect Evidence: Current Challenges. Insects. 2021;12(4):314.

2. Mendonça PM, dos Santos-Mallet JR, de Mello RP, Gomes L, Queiroz MM de C. Identification of fly eggs using scanning electron microscopy for forensic investigations. Micron. 2008;39(7):802–7.

3. Sukontason K, Sukontason KL, Piangjai S, Chaiwong T, Boonchu N, Kurahashi H, et al. Larval ultrastructure of Parasarcophaga dux (Thomson) (Diptera: Sarcophagidae). Micron. 2003 Dec 1;34(8):359–64.

4. Rivers DB, Dahlem GA. The Science of Forensic Entomology. Second Edition. John Wiley & Sons Ltd; 2023; p. 542.

5. Martoni F, Valenzuela I, Blacket MJ. Non-destructive DNA extractions from fly larvae (Diptera: Muscidae) enable molecular identification of species and enhance morphological features. Austral Entomology. 2019;58(4):848–56.

6. OSAC 2022-N-0039. Collecting & Preserving Entomological Evidence from a Terrestrial Environment Version 2.1 [Internet]. National Institute of Standards and Technology; 2025 [cited 2025 May 16]. Available from: https://www.nist.gov/system/files/documents/2024/04/23/OSAC%202022-N-0039%20Collecting%20%26%20Preserving%20Entomological%20Evidence%20from%20a%20Terrestrial%20Environment%20Version%202.0.pdf

7. Apasrawirote D, Boonchai P, Muneesawang P, Nakhonkam W, Bunchu N. Assessment of deep convolutional neural network models for species identification of forensically-important fly maggots based on images of posterior spiracles. Sci Rep. 2022;12(1):4753.

8. Sukontason KL, Sukontason K, Lertthamnongtham S, Kuntalue B, Thijuk N, Vogtsberger RC, et al. Surface Ultrastructure of Chrysomya rufifacies (Macquart) Larvae (Diptera: Calliphoridae). Journal of Medical Entomology. 2003;40(3):259–67.

9. Kress WJ, Erickson DL. DNA barcodes: Genes, genomics, and bioinformatics. Proc Natl Acad Sci U S A. 2008;105(8):2761–2.

10. Dong Z, Wang Y, Li C, Li L, Men X. Mitochondrial DNA as a Molecular Marker in Insect Ecology: Current Status and Future Prospects. Reddy GVP, editor. Annals of the Entomological Society of America. 2021;114(4):470–6.

11. Pimsler ML, Pape T, Johnston JS, Wharton RA, Parrott JJ, Restuccia D, et al. Structural and Genetic Investigation of the Egg and First-Instar Larva of an Egg-Laying Population of Blaesoxipha plinthopyga (Diptera: Sarcophagidae), a Species of Forensic Importance. Journal of Medical Entomology. 2014;51(6):1283–95.

12. Kumar V, Seal DR, Osborne LS, McKenzie CL. Coupling scanning electron microscopy with DNA bar coding: a novel approach for thrips identification. Appl Entomol Zool. 2014;49(3):403–9.

13. The Galaxy Community, Abueg LAL, Afgan E, Allart O, Awan AH, Bacon WA, et al. The Galaxy platform for accessible, reproducible, and collaborative data analyses: 2024 update. Nucleic Acids Research. 2024;52(W1):W83–94.

14. Huang X, Madan A. CAP3: A DNA Sequence Assembly Program. Genome Res. 1999;9(9):868–77.

15. Altschul SF, Gish W, Miller W, Myers EW, Lipman DJ. Basic Local Alignment Search Tool. Journal of Molecular Biology. 1990;215(3):403–10.

16. Benson DA, Cavanaugh M, Clark K, Karsch-Mizrachi I, Lipman DJ, Ostell J, et al. GenBank. Nucleic Acids Res. 2017;45(D1):D37–42.

17. Blankenberg D, Taylor J, Schenck I, He J, Zhang Y, Ghent M, et al. A framework for collaborative analysis of ENCODE data: Making large-scale analyses biologist-friendly. Genome Res. 2007;17(6):960–4.

18. Rice P, Longden I, Bleasby A. EMBOSS: The European Molecular Biology Open Software Suite. Trends in Genetics. 2000;16(6):276–7.

19. Larkin MA, Blackshields G, Brown NP, Chenna R, McGettigan PA, McWilliam H, et al. Clustal W and Clustal X version 2.0. Bioinformatics. 2007;23(21):2947–8.

20. Stothard P. The Sequence Manipulation Suite: JavaScript Programs for Analyzing and Formatting Protein and DNA Sequences. BioTechniques. 2000;28(6):1102–4.

21. Tardi NJ, Cook ME, Edwards KA. Rapid phenotypic analysis of uncoated Drosophila samples with low-vacuum scanning electron microscopy. Fly. 2012;6(3):184–92.

22. Thomsen PF, Elias S, Gilbert MTP, Haile J, Munch K, Kuzmina S, et al. Non-Destructive Sampling of Ancient Insect DNA. DeSalle R, editor. PLoS ONE. 2009;4(4):e5048.

23. Rusch TW, Adutwumwaah A, Beebe LEJ, Tomberlin JK, Tarone AM. The upper thermal tolerance of the secondary screwworm, Cochliomyia macellaria Fabricius (Diptera: Calliphoridae). Journal of Thermal Biology. 2019;85:102405.

24. Estes AM, Hearn DJ, Bronstein JL, Pierson EA. The Olive Fly Endosymbiont, “Candidatus Erwinia dacicola,” Switches from an Intracellular Existence to an Extracellular Existence during Host Insect Development. Applied and Environmental Microbiology. 2009;75(22):7097–106.

25. Mitchell A. Collecting in collections: a PCR strategy and primer set for DNA barcoding of decades-old dried museum specimens. Molecular Ecology Resources. 2015;15(5):1102–11.

26. Mehta A, Haber JE. Sources of DNA Double-Strand Breaks and Models of Recombinational DNA Repair. Cold Spring Harb Perspect Biol. 2014;6(9):a016428.

27. Fuyuhiko T, Yoshikawa K. The Enzymes. Volume 55. Elsevier; 2022; p. 1–5.

28. Sikorsky JA, Primerano DA, Fenger TW, Denvir J. Effect of DNA damage on PCR amplification efficiency with the relative threshold cycle method. Biochemical and Biophysical Research Communications. 2004;323(3):823–30.

29. Kranzfelder P, Ekrem T, Stur E. Trace DNA from insect skins: a comparison of five extraction protocols and direct PCR on chironomid pupal exuviae. Molecular Ecology Resources. 2016;16(1):353–63.

30. Altincicek B, Stötzel S, Wygrecka M, Preissner KT, Vilcinskas A. Host-Derived Extracellular Nucleic Acids Enhance Innate Immune Responses, Induce Coagulation, and Prolong Survival upon Infection in Insects. J Immunol. 2008;181(4):2705–12.

31. Boehme P, Amendt J, Zehner R. The use of COI barcodes for molecular identification of forensically important fly species in Germany. Parasitol Res. 2012;110(6):2325–32.

32. Stasiukynas L, Havelka J, da Silva FL, Torres Jimenez MF, Podėnas S, Lekoveckaitė A. COI Insights into Diversity and Species Delimitation of Immature Stages of Non-Biting Midges (Diptera: Chironomidae). Insects. 2025;16(2):174.

33. Chen H, Dong H, Yuan H, Shan W, Zhou Q, Li X, et al. Mitochondrial COI and Cytb gene as valid molecular identification marker of sandfly species (Diptera: Psychodidae) in China. Acta Tropica. 2023;238:106798.

34. Nelson LA, Wallman JF, Dowton M. Using COI barcodes to identify forensically and medically important blowflies. Medical and Veterinary Entomology. 2007;21(1):44–52.

35. Yusseff-Vanegas SZ, Agnarsson I. DNA-barcoding of forensically important blow flies (Diptera: Calliphoridae) in the Caribbean Region. PeerJ. 2017;5:e3516.

36. Richards CS, Crous KL, Villet MH. Models of development for blowfly sister species Chrysomya chloropyga and Chrysomya putoria. Medical & Veterinary Entomology. 2009;23(1):56–61.

