## Supplemental images for "Insect Identification with ESEM Imaging and DNA Barcoding for Forensic Use"

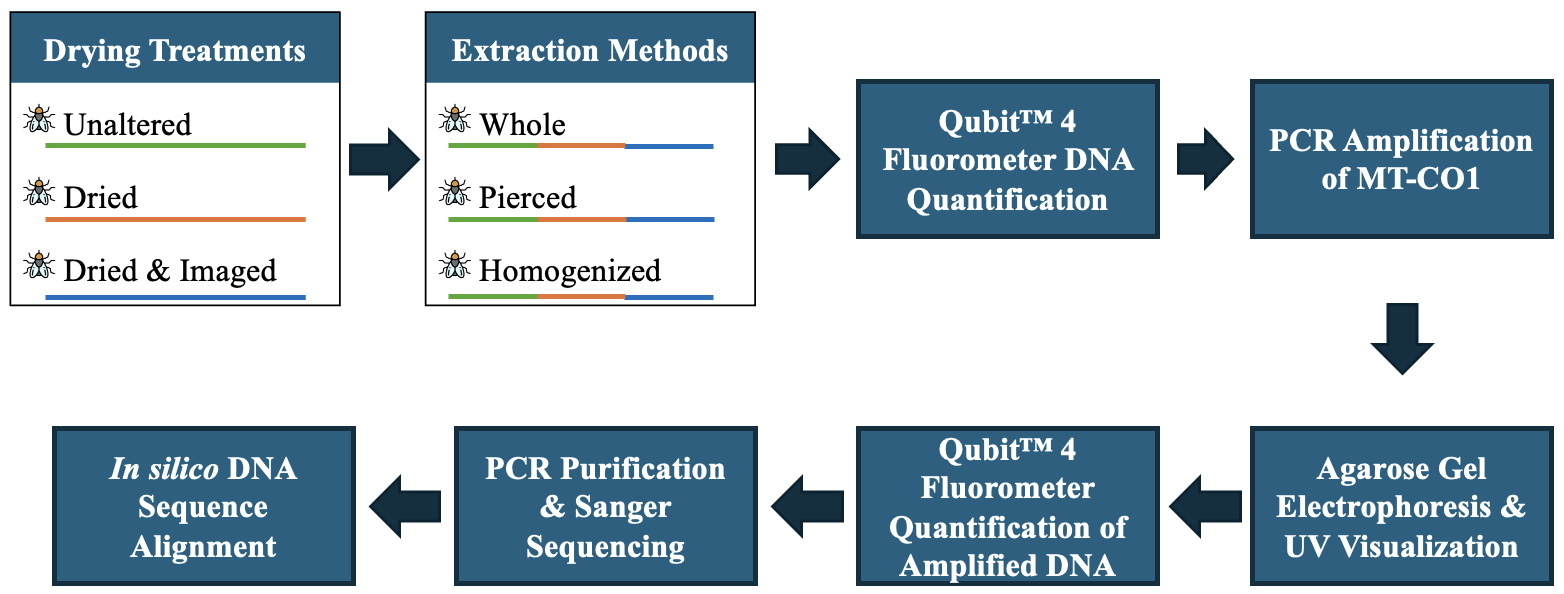
Supplemental images

**Supplemental Figure 1.** Overview of the experimental workflow, from specimen treatment DNA extraction to PCR amplification, sequencing, ending at data analysis.

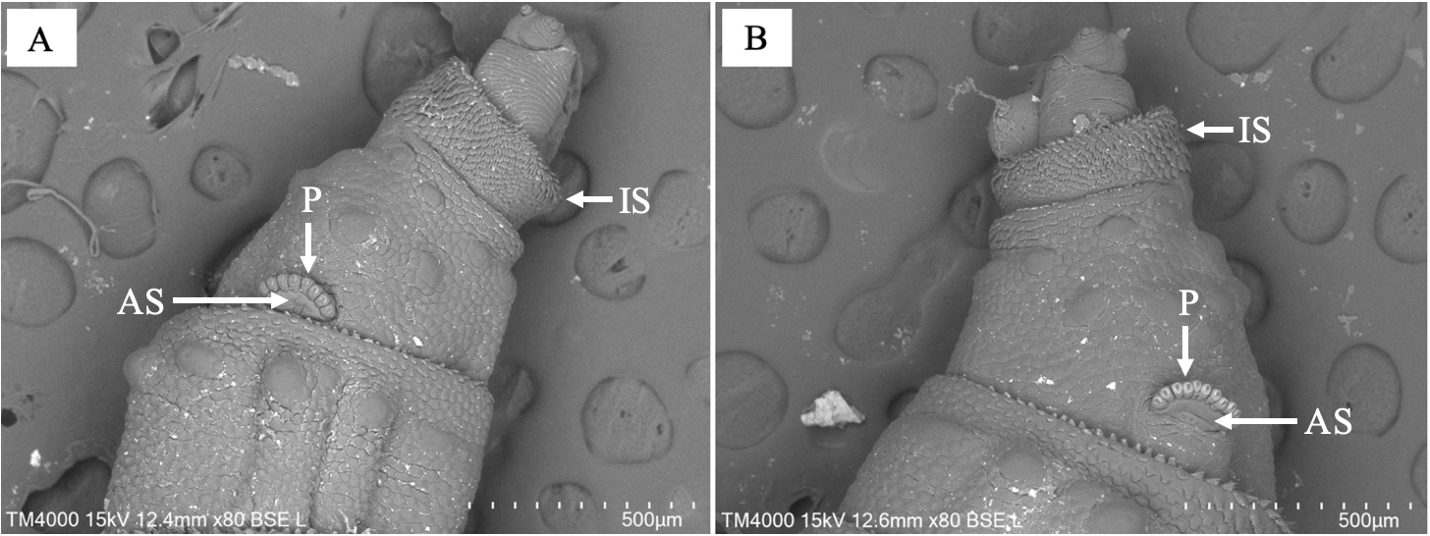

**Supplemental Figure 2.** Comparison of the anterior end of Chrysomya rufifacies larvae using ESEM at 80x magnification with BSE detection signal in left orientation: (A) dried larva sample and (B) undried larva sample.

**Supplemental Table 1.** Total DNA Yield (ng) post DNA extraction from different specimen treatments per extraction method.

| **Specimen Treatment** | **Extraction Method (ng)** | | |
| --- | --- | --- | --- |
|  | *Whole* | *Poked* | *Homogenized* |
| Untreated | 93.36 | 83.58 | 136.32 |
| Drying | 100.65 | 108.65 | 122.00 |
| ESEM | 74.02 | 91.34 | 112.35 |

**Supplemental Table 2.** Analysis of Variance (ANOVA) of DNA yield (ng/µL) from larval specimens post extraction using different extraction methods and specimen treatments.

| **Source of Variability** | **Degrees of Freedom** | **Sum of Squares** | **Mean Square** | **F Statistic** | **p-Value** |
| --- | --- | --- | --- | --- | --- |
| Extraction Method | 2 | 8.16 | 4.081 | 2.669 | 0.0754 |
| Specimen Treatment | 2 | 1.98 | 0.992 | 0.649 | 0.5255 |
| Interaction | 4 | 2.01 | 0.503 | 0.329 | 0.8576 |
| Residuals | 81 | 123.83 | 1.529 |  |  |

**Supplemental Table 4.** Mean of amplicon concentration (ng/µL) post PCR from different specimen treatments per extraction method.

| **Specimen Treatment** | **Extraction Method*** | | |
| --- | --- | --- | --- |
|  | *Whole* | *Poked* | *Homogenized* |
| Untreated | 2.9114 | 7.4500 | 11.6610 |
| Drying | 4.089 | 6.4900 | 12.9040 |
| ESEM | 4.3388 | 6.8820 | 12.7520 |
| **Mean** | 3.7797a | 6.9407b | 12.4390c |
| *Values are *highly significant; p < 0.05* | | | |

**Supplemental Table 5.** Analysis of Variance (ANOVA) of DNA concentration (ng/µL) from larval specimens post PCR using different extraction methods and specimen treatments.

| **Source of Variability** | **Degrees of Freedom** | **Sum of Squares** | **Mean Square** | **F Statistic** | **p-Value** |
| --- | --- | --- | --- | --- | --- |
| Extraction Method | 2 | 1152.1 | 576.0 | 65.204 | <2e^-16^* |
| Specimen Treatment | 2 | 6.9 | 3.4 | 0.388 | 0.679 |
| Interaction | 4 | 18.6 | 4.7 | 0.527 | 0.716 |
| Residuals | 81 | 715.6 | 8.8 |  |  |
